# A ketogenic diet mitigates hippocampal astrogliosis in epileptic brain

**DOI:** 10.1101/2025.04.11.647636

**Authors:** Jae Hyouk Choi, Hugo J. Kim, Shuhe Wang, Yueyang Cai, Ananya Achanta, Srija Pamujula, Hamza M. Thange, Matthew Shtrahman, Jong M. Rho

## Abstract

The ketogenic diet (KD) is an established treatment for patients with medically intractable epilepsy and is chiefly characterized by high fat/low-carbohydrate intake and the production of ketone bodies (KB) such as β-hydroxybutyrate (BHB). However, after more than a century of clinical use, the mechanisms underlying its efficacy remain unclear. While prior investigations have examined the effects of the KD and its metabolic substrates on synaptic transmission, few studies have explored a potential connection between astrocytic ion channels and seizure genesis. One essential function of astrocytes is spatial potassium buffering which influences passive potassium conductance (PPC), and when impaired, can result in neuronal hyperexcitability. In the present study, we demonstrate that the KD can mitigate hippocampal astrogliosis in the *Kcna1*-null (KO) mouse model of developmental epilepsy. Specifically, we observed a significant increase in GFAP expression in KO mice fed a control diet compared to wild-type (WT) mice, and that the KD prevented this change. Furthermore, we noted a reduction in hippocampal astrocytic PPC in epileptic mice, whereas KD-treated KO animals exhibited nearly normal passive conductance levels. In this regard, we found that while Kir4.1, TREK-1 and TWIK-1 RNA expression levels were not significantly altered by KD treatment in either WT or KO mice, BHB appeared to only minimally affect Kir4.1-mediated currents in transfected HEK cells. Furthermore, bulk RNA-seq analysis of the various treatment groups revealed KD-induced down-regulation of factors linked to hippocampal astrogliosis. Our findings indicate that the KD protects against epilepsy-associated astrogliosis and astrocytic PPC changes, underscoring a novel mechanism of action, and implicate extracellular potassium in its anti-seizure effects.

## Introduction

The ketogenic diet (KD) has been used to treat patients with epilepsy for over a century (Pluta & Jabłoński, 2011). This dietary regimen is characterized by high-fat and low-carbohydrate intake (Rusek et al., 2019) which induces a systemic transition from glycolytic to intermediary metabolism wherein fatty acids are oxidized and result in the production of ketone bodies (KB) such as acetoacetate and β-hydroxybutyrate (BHB) as energy substrates and signaling molecules (Düking et al., 2022). Importantly, the KD has been shown to block spontaneous recurrent seizures (SRS) in the approximately one-third of patients with epilepsy who fail to respond to anti-seizure medications or ASMs (Neal et al., 2008; T. A. Simeone et al., 2017; Ułamek-Kozioł et al., 2019). In parallel, there is substantial evidence in animal models for the anti-seizure efficacy of the KD (D’Andrea Meira, Romão, Pires Do Prado, et al., 2019; Rho, 2017; Rho & Boison, 2022; Simeone et al., 2017, 2021; Stafstrom & Rho, 2012).

Despite the advent of many new ASMs with notable improvements in pharmacokinetic profiles, the clinical efficacy rates remain largely unchanged (Chen et al., 2018). One potential limitation may be that prior mechanistic and experimental therapeutic investigations have primarily focused on neurons and less on glia and neuron-glia interactions. Recently, astrocytic ion channels have been implicated in seizure genesis in both animal models and surgically-resected human epileptic tissues; specifically, a reduction in the expression of Kir4.1 channels (encoded by either *Kcnj10* or *KCNJ10 genes*) has been reported (Bedner et al., 2015; Song et al., 2018; Tong et al., 2014). A principal function of astrocytes is potassium buffering – i.e., the passive uptake/release of the potassium ions to maintain ionic homeostasis, which is essential for normal neuronal activity (Zhou et al., 2009; Mi Hwang et al., 2014; Bae et al., 2020). Impairment of potassium buffering results in an increase in the extracellular potassium concentration, which then provokes neuronal excitability (Kelley et al., 2018; Zhong et al., 2023; Zhou et al., 2009)

There is now broad recognition that most if not all brain pathologies can result in part from dysfunction of both neurons and glial cells (Lee et al., 2022). Astrocytes play crucial roles in maintaining a healthy brain, including the regulation of neurotransmitter balance and ionic homeostasis (Volterra & Meldolesi, 2005). The term “reactive astrogliosis” encompasses the changes in astrocyte morphology, biochemistry, and physiology in response to damage in the central nervous system (CNS) (Sofroniew & Vinters, 2010). Astrogliosis associated with CNS injury and disease has been acknowledged for over a century (Escartin et al., 2021).

Recent advances have revealed that astrogliosis is a multifaceted process, encompassing a spectrum from subtle and reversible changes in gene expression and morphology to pronounced and long-lasting alterations linked with scar formation (Escartin et al., 2021). Relevant to the present study, astrogliosis is thought to contribute to the development and progression of epilepsy (Olsen & Sontheimer, 2008; Robel et al., 2015). In animal models, reactive astrocytes experience significant physiological changes that disrupt extracellular ion concentrations such as potassium, neurotransmitter balance, as well as intracellular and extracellular water levels. These alterations can lead to neuronal damage (Bordey et al., 2001; Bordey & Sontheimer, 1998). Together, multiple observations strongly suggest that decreased potassium buffering might be involved in the pathophysiology of epilepsy (Çarçak et al., 2023; Coulter & Steinhauser, 2015). Indeed, previous studies indicate that astrogliosis can be linked to several CNS disorders and to the involvement of numerous ion channels (Olsen et al., 2015). Interestingly, the Kir4.1 channel, a major contributor to the passive conductance of astrocytes, has been reported to be dysfunctional in many disease models (Djukic et al., 2007; Kelley et al., 2018; Mukai et al., 2018; Olsen & Sontheimer, 2008; Song et al., 2018; Tong et al., 2014).

Here, we examined whether the KD may mitigate astrogliosis in the epileptic brain and reinstate the functionality of ion channels responsible for normal astrocytic potassium buffering and passive membrane conductance. We further explored whether BHB can regulate Kir4.1 astrocytic ion channels to enhance potassium buffering. Finally, using bulk RNA-seq analysis, we examined the gene expression profiles of WT and KO mice fed either a KD for two weeks or SD to identify potential pathways and targets involved in the pathogenesis of aberrant gliosis and which might be amenable to therapeutic intervention.

## Results

### Ketogenic Diet Induces Anti-Seizure Effects in KO mice

We performed video-EEG recordings on KO and WT mice and compared the daily occurrence of seizures (see Methods below) with either standard diet (SD) or an experimental KD. Daily seizure counts significantly decreased in KO mice after two weeks of KD treatment compared to the SD-fed animals (*p=0.047*, n=3-4 per group) (Figure 1A, C, D), consistent with our previous observations (Kim et al., 2015; K. A. Simeone et al., 2016, 2021). Further, we confirmed that KD administration significantly increased blood BHB levels and reduced blood glucose levels without affecting body weight (Figure 1B). We and others have shown that BHB alone can render anti-seizure effects in various model systems through a multiplicity of mechanisms (Simeone et al., 2018) and can result in a higher ratio of gamma-aminobutyric acid (GABA) to glutamate, longer seizure latency, less severe SRS (Olson et al., 2018; Qiao et al., 2024), Collectively, these findings provide further support for the anti-seizure effects of the KD in rodent models of epilepsy.

**Figure 1.**
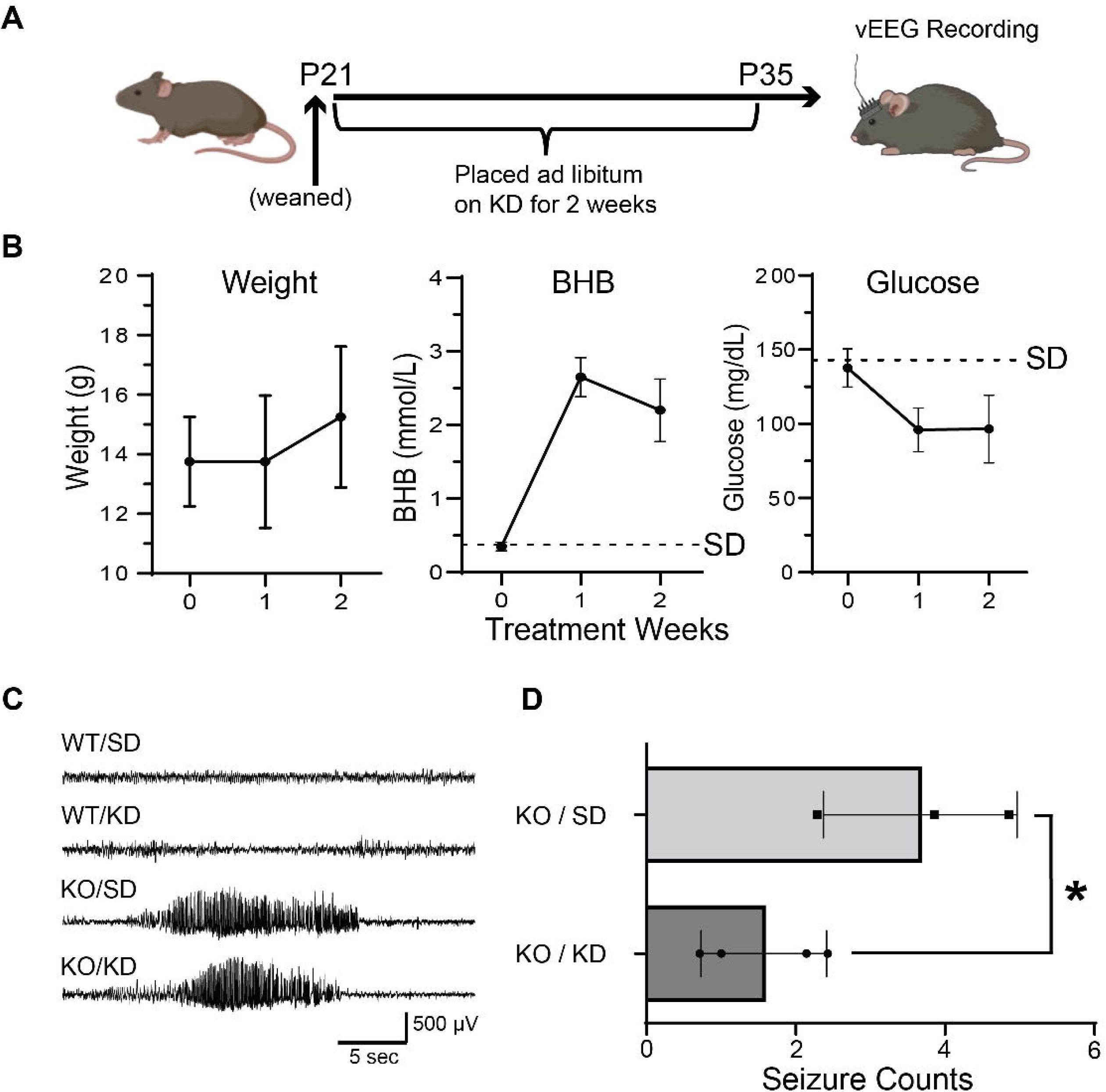
A ketogenic diet (KD) blocks spontaneous recurrent seizures and enhances EEG gamma power in epileptic mice. (A) Daily Seizure count significantly decreased after two weeks of KD treatment. (B) Representative tracings of EEG signals recorded in *Kcna1-*null (KO) mice and WT animals fed either (standard diet) SD or KD. For KO mice, epileptic events were similar as depicted. (C) Plasma β-hydroxybutyrate (BHB) and glucose levels, and weight changes were determined in KO mice treated with KD. Each symbol represents the mean ± S.E.M. BHB and glucose levels were also measured in mice treated with SD, as shown by the dotted line. (D) Relative gamma oscillation power was increased by the KD in KO mice. *Student t-test: *p*<0.05. (E) Schematic timeline for the vEEG recordings during dietary treatment.

### KD Can Reverse Gene Expression Changes in Hippocampus of Epileptic Mice

To gain insights into the mechanisms underlying the restorative effects of the KD in our KO mice, we performed bulk RNA-seq on hippocampi collected from WT and KO mice treated with a SD or KD. We anticipated that the varying experimental conditions might result in differentially expressed (DE) genes associated with glial cell responses in the epileptic brain. Between WT/SD and WT/KD groups, we identified 67 DE genes (False discovery rate (FDR) corrected *p* value < 0.05, |log2 fold change [fold change (FC)]| > 1) (Figure 2A). Notably, no significant biological processes were enriched in this gene set according to Gene Ontology (GO) analysis. This indicates that the effect of KD is minimal in WT mice. We next compared WT/SD and KO/SD groups, and found 332 DE genes, with most of these genes being overexpressed in KO mice fed a SD (Figure 2B). GO analysis revealed that pathways related to synaptic signaling were highly enriched among these DE genes. When we examined both WT and KO animals fed a KD, only 32 DE genes could be identified between WT/KD and KO/KD mice, and within these DE genes, only small subsets appeared overlapping between the two groups (Figure 2C).

**Figure 2.**
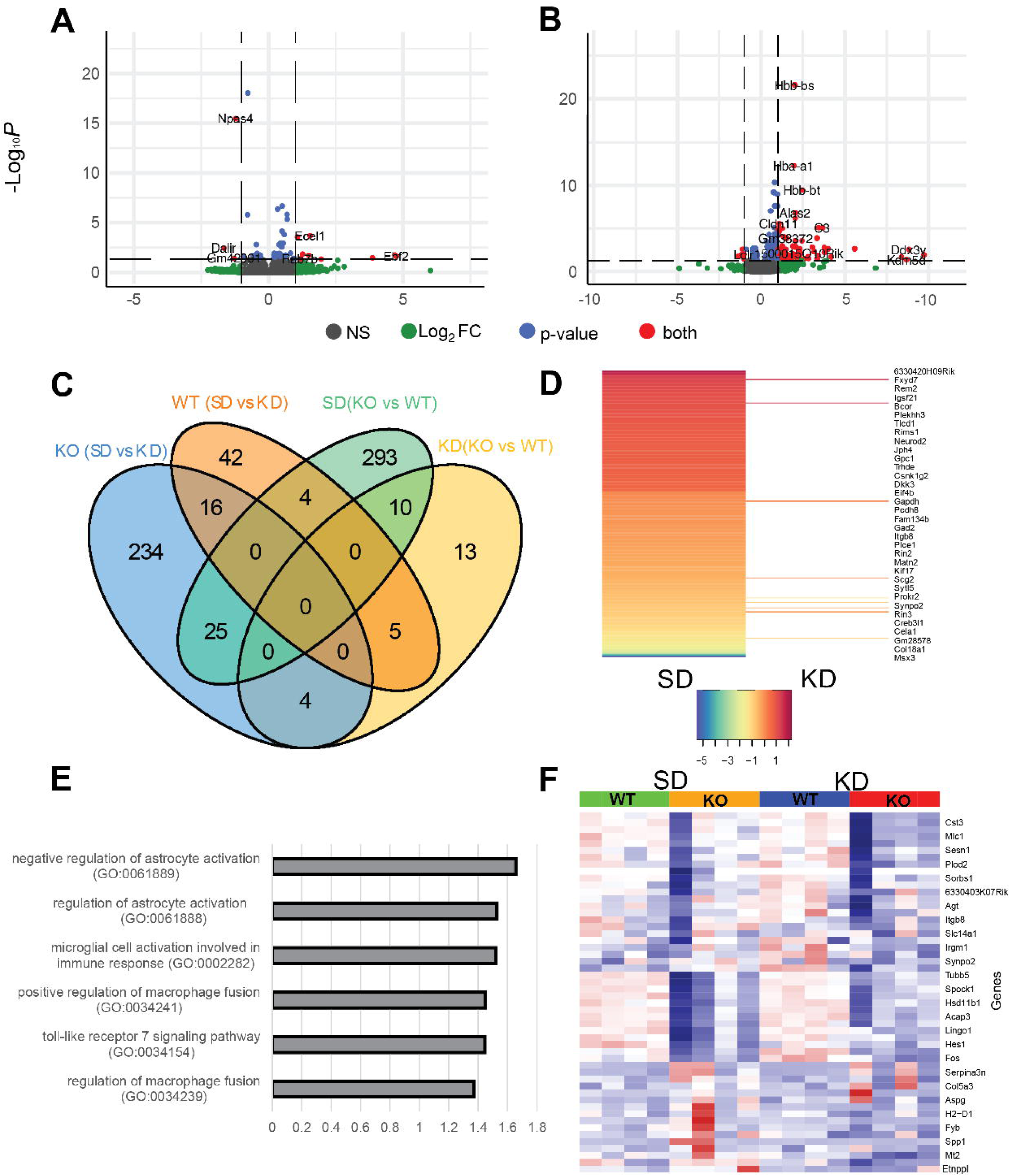
Bulk RNA-seq analysis of ketogenic diet effects in control and epileptic *Kcna1*- null (KO) mice. (A) Volcano plot between wildtype (WT) standard diet (SD) vs. ketogenic diet (KD) indicating that there are no major changes induced by the KD in control animals. (B) Volcano plot between KO/SD and KO/KD groups indicating that the KD can significantly change expression profiles in epileptic mice. (C) Number of differentially expressed (DE) genes between the four comparison groups (WT/SD, KO/SD, WT/KD, KO/KD) demonstrating the large number of DE genes between WT and KO animals fed either a SD or KD. (D) A significant number of DE genes identified in the SD condition are no longer significantly differentially expressed in the KD treatment group (adjusted *p*<0.05). (E) Gene Ontology terms for the Top 100 DE genes in the SD group no longer differentially expressed after KD treatment was enriched in pathways related to astrocyte activation. (F) Heatmap of genes regulating astrogliosis from the known literature (Matusova et al., 2023) of the four groups indicates relatively small changes in patterns when the KD is compared to the SD.

As earlier studies have shown that a four-way comparison can identify differentially regulated gene expression changes (Burda et al., 2022), we compared the DE genes between WT and KO mice fed either a SD or KD. Most of the DE genes found with SD treatment were no longer differentially expressed with the KD, indicating that the KD can mitigate the prominent gene expression changes seen in KO mice compared to WT controls (Figure 2D, Table S1). Biologically important changes are listed in Table 1. Also, the GO analysis of these differentially regulated genes showed higher enrichment in the regulation of the astrocyte pathway (GO: 0061888, 0061889) (Figure 2E). This result is consistent with a homeostatic effect of the KD in normalizing gene expression changes induced by astrocyte activation, and underscore the thesis that astrogliosis may be an important pathophysiological mechanism underlying the prominent epileptic phenotype in mice lacking the *Kcna1* gene.

Using a specific gene list related to astrogliosis (Matusova et al., 2023), we compared gene expression patterns across our four experimental groups: WT/SD, WT/KD, KO/SD, and KO/KD (Figure 2F). Our analysis revealed two distinct gene expression patterns. The first group comprised genes highly expressed in WT animals but were significantly decreased in KO mice and were consistent across both dietary conditions. By contrast, the second group exhibited high expression in KO animals and reduced expression in WT mice. Notably, the overexpression of these astrogliosis-regulating genes in KO animals was most pronounced during SD treatment. Critically, KD treatment appeared to attenuate the overexpression of astrogliosis-related genes in epileptic KO mice.

### The KD Mitigates Astrogliosis in KO Mice

Previous studies have reported significant alterations in the morphology and function of glial cells in various forms of epilepsy in both human surgical specimens and animal models, and such evidence indicates that astrocytes are likely involved in the pathogenesis of seizures and epilepsy (Fellin & Haydon, 2005). Specifically, there is growing evidence that epilepsy may arise from astrocytic pathology, as direct stimulation of these cells can lead to prolonged neuronal depolarization and epileptiform discharges (Chan et al., 2019; Hayatdavoudi et al., 2022; Verhoog et al., 2020). As stated above, bulk RNA-seq analysis (Figure 2) revealed that the KD reversed the overexpression of astrogliosis-related genes. To confirm whether these transcriptional changes corresponded to morphological hallmarks of reactive astrocytes, we performed semi-quantitative Immunohistochemistry (IHC) in KO mice before and after KD treatment. Interestingly, in our KO model, we observed significant hypertrophy of astrocytes (Figure 3A) in the hippocampus of KO mice (*p=0.0001*, KO/SD and WT/SD, n=7 each group). GFAP labeling was quantified using IHC data (Figure 3B) and was expressed as a percentage of the overall 2-D area in each standard field of view. Surprisingly, two weeks of KD treatment prevented aberrant astrogliosis and resulted in a level comparable to that seen in the WT/SD group (*p=0.40*, KO/KD compared to WT/KD, n=7 for each group) (Figure 3A-C, S1A). We next confirmed hippocampal protein levels of GFAP using western blots. GFAP protein expression was significantly increased in KO mice compared to controls, both fed a SD (*p*=0.0016, n=4 in each group). However, KD treatment in KO animals led to a reduction in GFAP protein levels comparable to that seen in the WT/KD group (*p=0.15*, n=4 in each group).

**Figure 3.**
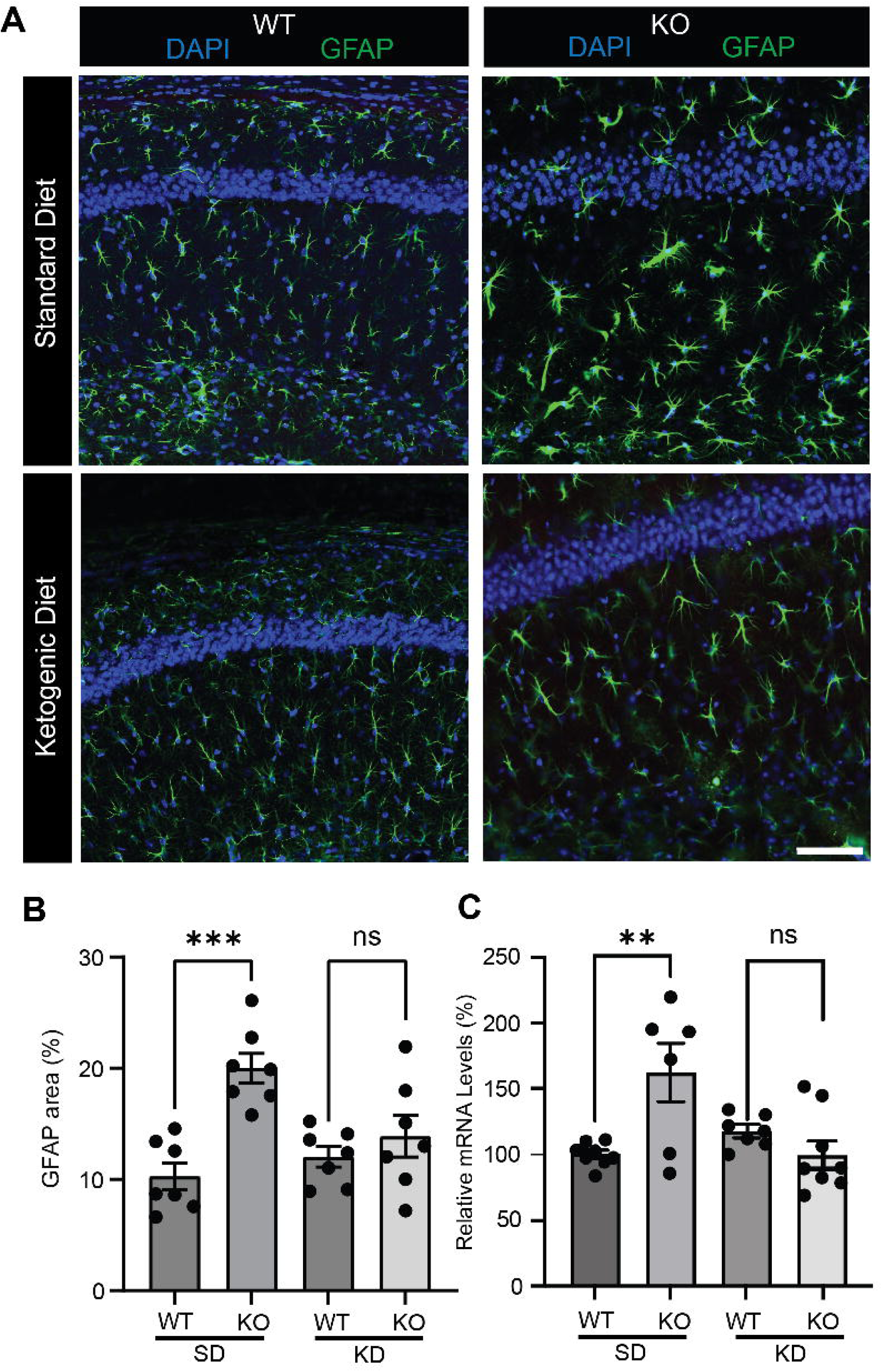
Representative images of GFAP immunoreactivity in CA1 hippocampal astrocytes. 10x magnification, scale bar 200 µm. (A) Astrocytes exhibited significant astrogliosis in *Kcna1*-null (KO) mice on a standard diet (SD) compared to wild-type (WT) controls on a SD. However, the aberrant astrogliosis was mitigated in the KO group after two weeks of ketogenic diet (KD) treatment. (B) Summary bar graphs showing the percentage of GFAP labeling as a percentage of total area in the field of view. *p* values were obtained with the Student’s t-test. \*\*\**p*<0.0001. (C) qRT-PCR analysis revealed a significant increase in GFAP RNA expression and that of another astrocytic marker, Kir4.1, in KO mice. However, KD administration for 2 weeks led to normalization of RNA expression levels. Expression levels of TWIK1 and TREK1 ion channels were also measured to assess for possible contributory effects to KD action, but these channels were not differentially expressed between the WT and KO groups.

Since altered ion channel function can be seen in reactive astrocytes, we asked whether we could detect either expression and/or functional changes in ion channels involved in passive potassium conductance. Specifically, we examined the mRNA expression of *Kir4.1* (*Kcnj10*), *TWIK-1* (*Kcnk1*), and *TREK-1* (*Kcnk2*) channels because they are known to contribute to astrocytic spatial potassium buffering (Bae et al., 2020; Mi Hwang et al., 2014; Zhou et al., 2009). We found that Kir4.1 channel mRNA expression was increased in the hippocampus of our KO mice (Figure S1B; *p=0.0075,* KO/SD compared to WT/SD groups, n=8 and 6, respectively). Given this finding, we calculated *Kir4.1* expression normalized to GFAP expression area (divided by volume) in hippocampus and found that Kir4.1 was decreased in KO mice compared to WT controls but was restored by KD treatment (Figure S1C). However, there were no significant changes in hippocampal mRNA levels of *TWIK-1* and *TREK-1* between KO mice versus WT controls (Figure S1D).

### The KD Rescues Astrocytic Passive Conductance Changes in KO Mice

Glial cells play diverse roles in the brain, including regulating nutrient supply, transmitting signals from the blood to neurons, clearing synaptic glutamate, guiding neuronal growth, and managing the distribution of potassium (K^+^) ions. The long-range spatial buffering of K^+^ by glial cells serves as a mechanism for synchronizing or spreading activity during paroxysmal oscillations. Glial cells also release neuroactive substances and modulate synaptic transmission by altering ion channels, gap junctions, receptors, and transporters (Djukic et al., 2007; Olsen & Sontheimer, 2008). As mentioned above, altered astrocytes in the epileptic mouse brain could induce neuronal hyperexcitability (Verhoog et al., 2020) and potentially contribute to seizure genesis (and possibly, epileptogenesis).

Furthermore, because abnormally increased neuronal firing in the epileptic brain can generate more potassium in the extracellular space, the role of astrocytes in spatial potassium buffering is critical toward restoring normal cellular membrane potentials (Mukai et al., 2018; Song et al., 2018). Therefore, we examined whether the PPC of CA1 hippocampal astrocytes is altered in our epileptic KO mice using whole-cell patch clamp recording techniques. We found that the passive conductance was significantly decreased in hippocampal astrocytes from KO mice compared to WT controls (Figure 4A-B, S2). The inward rectifying currents were significantly decreased in the KO*/*SD group versus WT/SD animals (*p=0.0077,* n=6 and 7, respectively) (Figure 4C-D, S2B). In addition, although the outward rectifying current was decreased in KO mice compared to WT controls on a SD (*p=0.0005,* n=6 and 7, respectively), KD treatment restored this current to the normal range of passive conductance (inward current: *p=0.066*, outward current: *p=0.60*, KO/KD vs. WT/KD; n=5 in each group) (Figure 4A-D, S2C). Additionally, as dysfunction in astrocytic ion channels has been linked to increased neuronal firing (Verhoog et al., 2020), we measured the rheobase current to further evaluate the excitability of CA1 neurons adjacent to hippocampal astrocytes (Figure 4E, S2-3). The firing frequency of CA1 pyramidal neurons in KO mice was significantly decreased after treatment with the KD (Figure 4F-G), likely due to restoration of the PPC.

**Figure 4.**
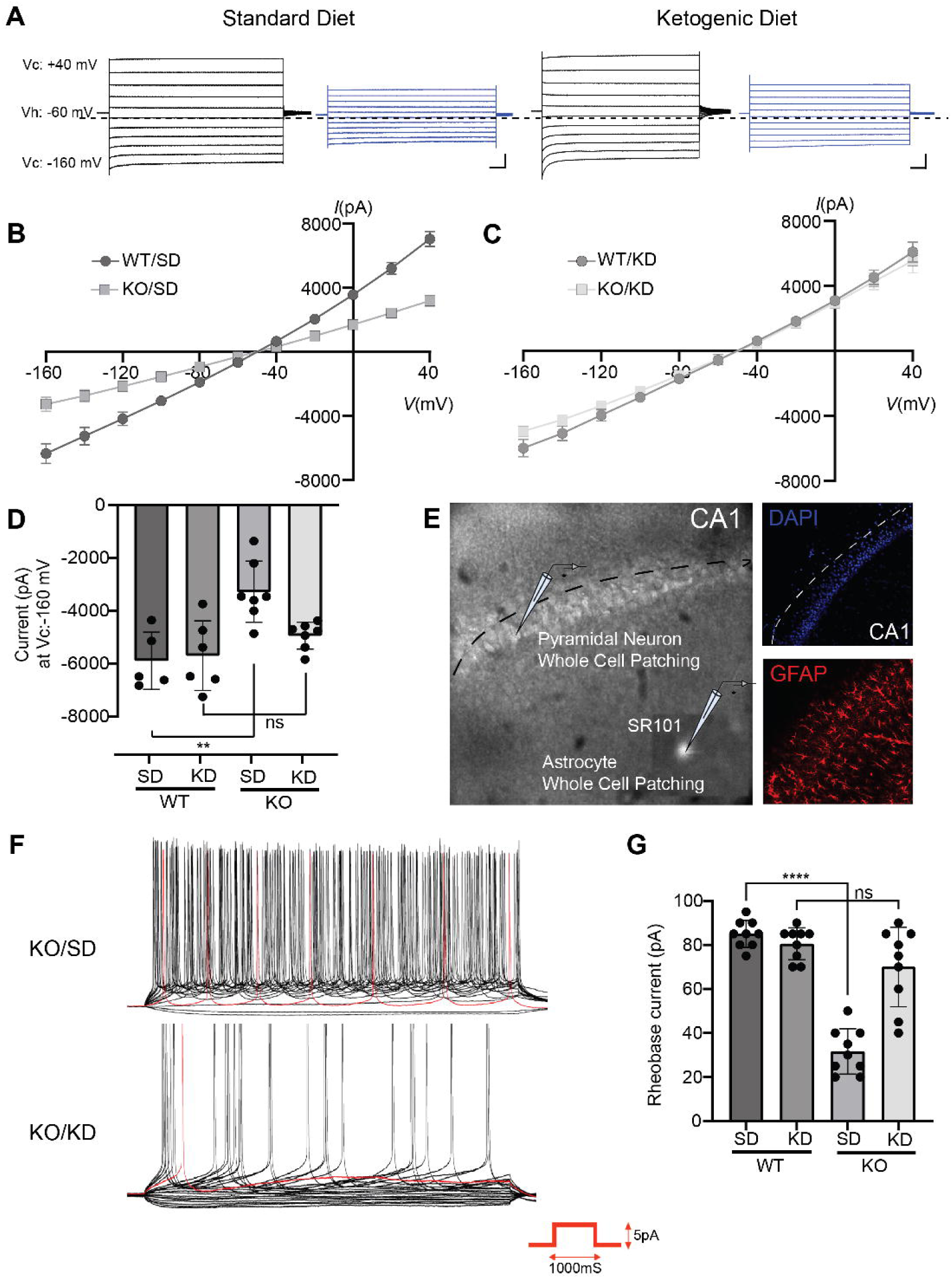
Whole-cell currents measured in hippocampal astrocytes and CA1 pyramidal neurons of wild-type (WT) and *Kcna1*-null (KO) mice on either standard diet (SD) or ketogenic diet (KD). (A) Representative traces reflecting passive conductance levels from hippocampal astrocytes on SD/KD. Using a command voltage pulse protocol, voltages were stepped from −160 mV to +40 mV in increments of 20 mV with a holding potential of −60 mV. (B-C) The current–voltage relationship (I–V) was assessed from −160 mV to +60 mV (B) Bar graph summary of inward currents; this gap shows the inward current levels between WT vs. KO mice fed either SD or KD: *p* values (SD:0.0016, KD:0.096). (D) Bar graph of current levels at −160 mV, indicating inward currents in hippocampal astrocytes from WT or KO mice fed either SD or KD. (E) A low power view of the hippocampus using infrared differential interference contrast (IR-DIC) microscopy. Recordings were made in acute brain slices with patch electrodes in CA1 pyramidal cells and hippocampal astrocytes. Also depicted are images of DAPI and GFAP immunoreactivity in the CA1 region using the same hippocampal slices. (F-G) Representative traces of the membrane potential to stepwise current injections recorded from CA1 pyramidal neurons of KO mice compared to W controls on either a SD (n=9) or a KD (n=5). The resting membrane potential remained at −70 mV. Depolarizing currents were injected in increments of 5 pA. CA1 neurons in KO mice (n=5) were more excitable compared to WT mice (n=9) while both were fed a SD (\*\*\*\**p*<0.0001). However, this difference was diminished after KD treatment; KO/KD compared to WT/KD, *p*=0.12. Data represent the mean ± S.E.M., Student’s t-test.

In summary, we discovered significant astrogliosis and increased GFAP expression in the hippocampus of *Kcna1*-null (KO) mice, and that two weeks of KD administration led to an amelioration of astrogliosis, improvement in astrocytic PPC and increases in Kir4.1 expression – the latter effects reflecting rescue of normal astrocytic ion channel function.

### D-BHB Tends to Enhance Passive Potassium Conductance in Kir4.1-Transfected HEK Cells

Since KB such as acetoacetate (AcAc) and BHB are synthesized by the liver in response to a high-fat KD, and to reduced levels of blood glucose and insulin (Cuenoud et al., 2020), we asked whether BHB (the primary substrate elaborated by the KD and one that can directly induce anti-seizure effects (Simeone et al., 2018) might exert modulatory activity on Kir4.1 channels transfected in HEK cells. We measured the passive conductance of Kir4.1 and compared this finding in control GFP vector-transfected HEK cells (Figure 5A-B). We observed an increase in inward rectifying currents in Kir4.1-GFP compared to control-GFP cells after 10 min of BHB (5 mM) bath application. However, at the tested concentration, BHB did not significantly alter the baseline Kir4.1 current (Figure 5C-D; *p*=0.14, n=15), and similarly, BHB treatment did not affect the inward rectifying current in control-GFP cells (Figure 5C; *p*=0.92, n=8).

**Figure 5.**
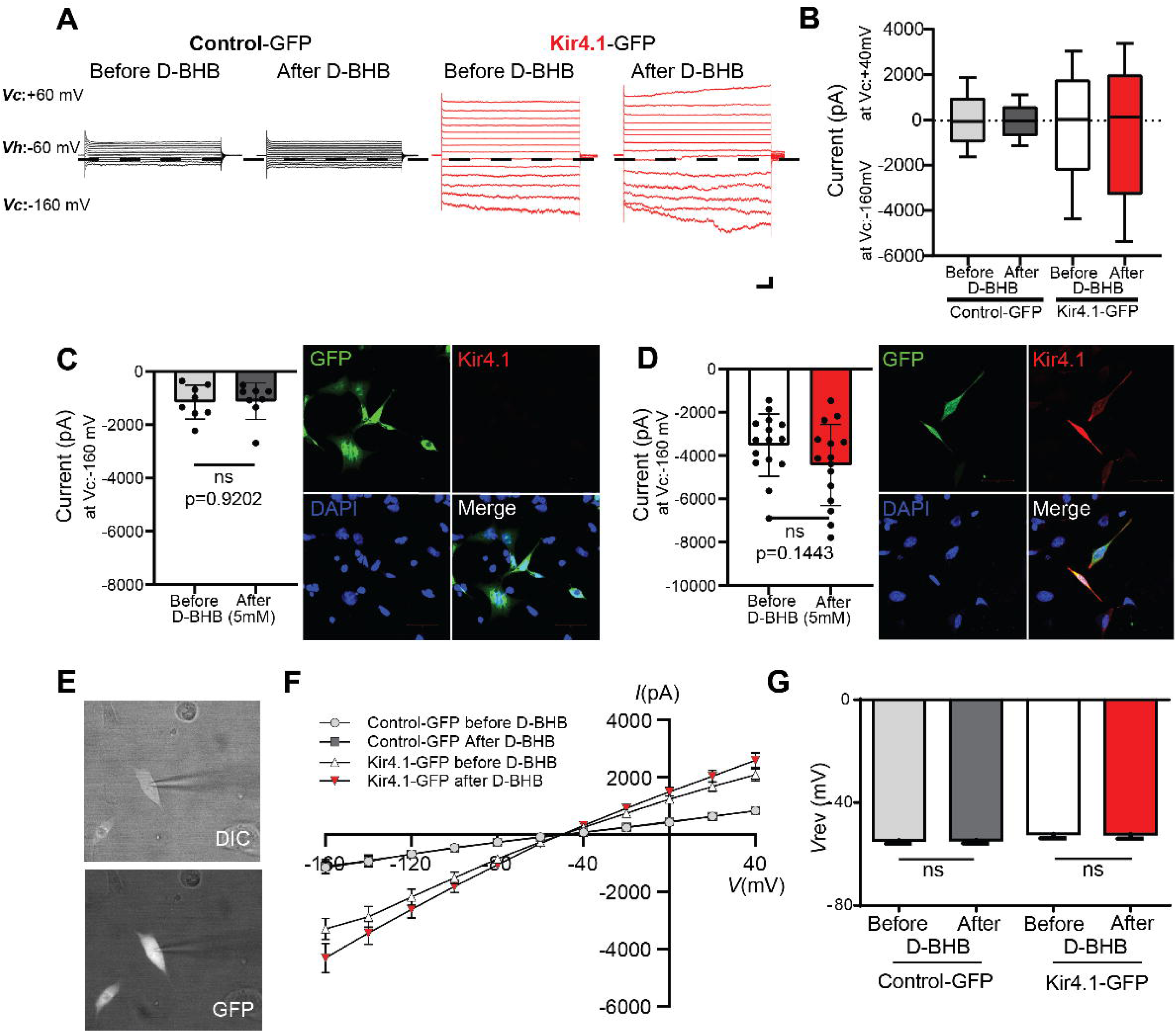
BHB Enhances Kir4.1 Currents After Bath Application. (A) Whole-cell currents in HEK cells transfected with control-GFP (left) or Kir4.1-GFP (right) before and after bath application of BHB (5 mM) for 10 min. Command voltage pulse protocol: voltages were stepped from −160 mV to +40 mV in increments of 20 mV from a holding potential of −60 mV. (B) Summary data; the dotted line indicates 0 pA. (C) Representative immunohistochemical images for Kir4.1-transfected HEK cells and summary bar graphs for inward currents Control- GFP (*p* value :0.9202) and Kir4.1-GFP (*p* value :0.1443). (E) DIC (differential interference contrast) (upper) and fluorescence (lower) images of HEK cells transfected with Control-GFP and Kir4.1-GFP. The control and Kir4.1 constructs also expressed a green fluorescence signal. (F) Representative whole-cell I–V curves in control-GFP and Kir4.1-GFP before and after BHB bath application. The series resistance was <15 MΩ, which we monitored throughout experiments. (G) The bar graph representing reversal potential before and after BHB. Data are mean ± S.E.M. *p* values were obtained with Student’s t-test. \**p*<0.05, \*\**p*<0.01.

Transfected Kir4.1 and control-GFP cells showed a current–voltage (I–V) relationship with a complex shape, consisting of both an outwardly rectifying outward current and a weakly inwardly rectifying inward current (Figure 5E-F). However, there was no significant shift in the reversal potential before and after BHB bath application: control-GFP, -54.73 mV±1.056 after BHB: -54.58±-1.28, Kir4.1-GFP, before BHB, -52.17±1.71, after D-BHB, -52.2±1.81 (Figure 5G).

These results suggest that BHB could potentially activate the gating mechanism of the *Kir4.1* channel. This finding aligns with our group’s previous report, indicating that KB can modulate neuronal firing by interacting with ATP-sensitive potassium (K_ATP_) channels (Kim et al., 2015), Indeed, the Kir4.1 channel is recognized as an ATP-sensitive inward rectifier potassium channel channel (Kubo et al., 2005).

### KD-Induced Mitigation of Astrogliosis Reveals Transcriptional Changes with Broader Neurological Implications

Considering that genes involved with astrocyte activation were less expressed under conditions of KD treatment, it is reasonable to expect a significant recovery of astrocytic functions such as potassium ion buffering after this dietary intervention. Furthermore, astrogliosis can induce hyperexcitability in neighboring CA1 neurons by impairing spatial potassium buffering in the epileptic brain. The KD may restore normal astrocytic morphology and rescue PPC function under these conditions (Figure 6A). Some of these changes were preventive while others could be involved in rescue. Henceforth, the gene expression changes in KO mice and those seen with KD rescue will be referred to as the ‘epilepsy factors’ and ‘ketogenic factors’, respectively.

**Figure 6.**
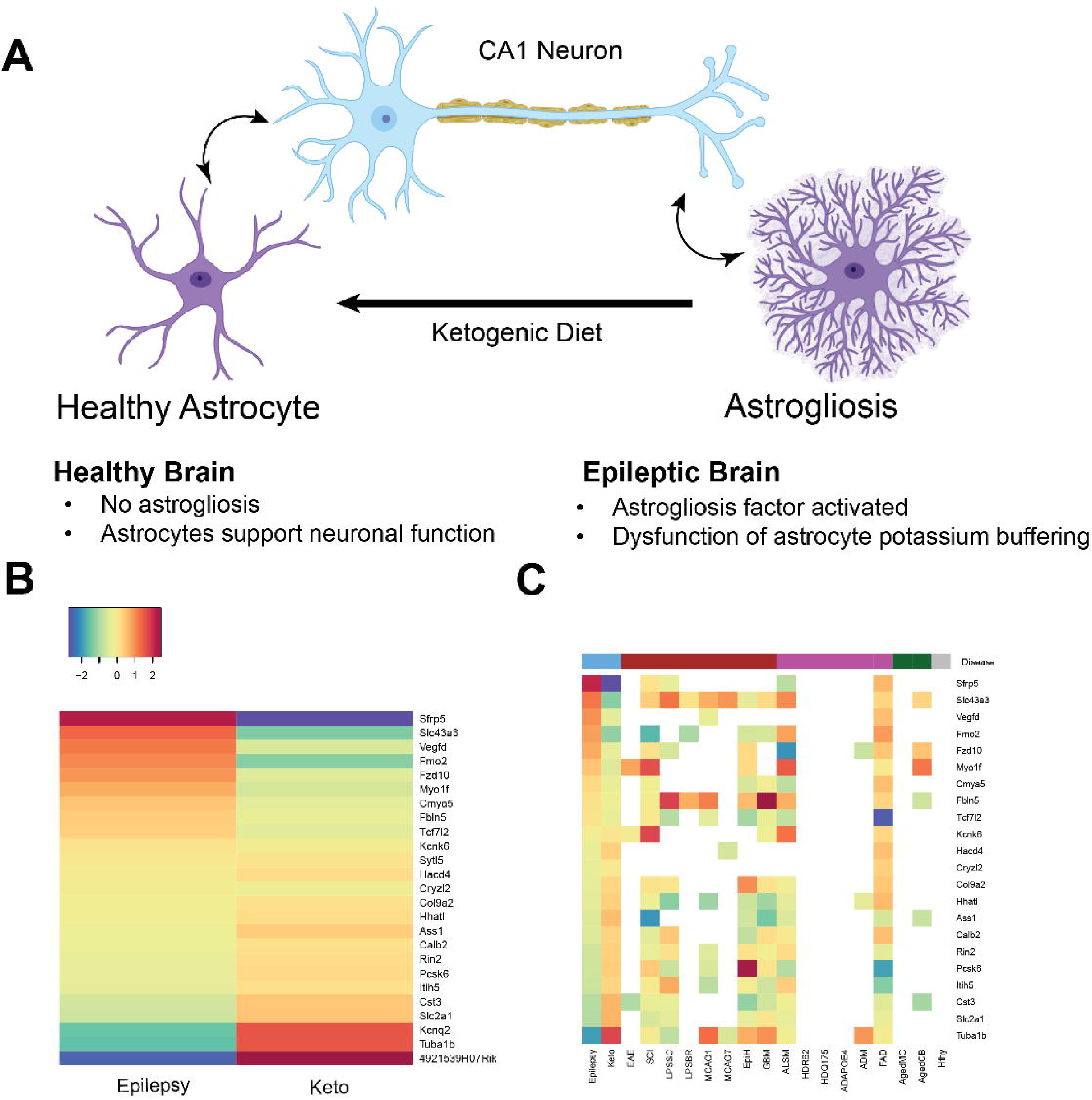
The potential effect of Ketogenic Diet in neurological disorders. (A) Astrogliosis factors are mitigated in the hippocampus of the epileptic brain after two weeks of a ketogenic diet (KD), and aberrant astrocytic potassium buffering is reversed. (B) List of genes reversed with KD treatment. (C) Gene expression profiles of reversed gene in other disease states listed in the Astrocyte Reactivity Regulation Browser (http://tr.astrocytereactivity.com/home).

Using our bulk RNA-seq data, we examined epilepsy factors by comparing the WT/SD and KO/SD groups and ketogenic factors by comparing KO/SD and KO/KD cohorts. Out of 332 DE genes between WT/SD and KO/SD groups, 26 DE genes were reversibly expressed between KO/SD and KO/KD groups (Figure 6B, Table S2). Three genes (*4921539H07Rik*, *Tuba1b*, *Kcnq2*) that are significantly downregulated by epilepsy factors were significantly upregulated by ketogenic factors. The other 23 genes were upregulated by epilepsy factors and downregulated by ketogenic factors. These patterns indicate that there are significant diet-induced reversals of gene expression profiles in the epileptic brain.

The pathways that regulate gene expression are shared to varying degrees across multiple neurological disorders and in the aging processes (Burda et al., 2022). As such, it is reasonable to speculate that the KD-induced effects on gene expression profiles could be applicable to conditions outside of epilepsy such as spinal cord injury (SCI) and traumatic brain injury (TBI) (Yarar-Fisher et al., 2021). In this light, we examined the reported changes in gene expression patterns described by Burda et al, (2022), including the following: experimental allergic encephalomyelitis (EAE) mouse, mouse model of SCI, lipopolysaccharide (LPS)-induced inflammation in spinal cord and brain of mice (LPSSC and LPSBR), ischemic reperfusion on days 1 and 7 in mice (MCAO1, MCAO7), human epilepsy (EpiH), human glioblastoma (GBM), mouse amyotrophic lateral sclerosis models (ALSM), two Huntington’s Disease mouse models (HDR62, HDQ175), human Alzheimer’s disease (ADAPOE4), and two mouse Alzheimer’s disease models (ADM, FAD), aged (2 years) motor cortex and cerebellum (AgedMC, AgedCB), and healthy (Hty) (Figure 6C).

Using comparison data normalized to the healthy dataset, we found a similar pattern in many of these disorders. The mouse Alzheimer’s disease model FAD correlated with our epilepsy model (R=0.34). We also found that the directionality of many gene expression changes was the same compared to the SCI and ALM models. Through these comparisons, we found correlations with some of the genes that were differentially expressed between WT and KO conditions and pathologic changes being reversed with KD treatment.

## Discussion

The KD has been utilized successfully in patients with medically intractable epilepsy for over a century, but the mechanisms underlying its clinical benefits have remained elusive. While the KD has predominantly been examined in the context of synaptic transmission (Düking et al., 2022; Fedorovich et al., 2018; Gom et al., 2021; T. A. Simeone et al., 2017; Spigoni et al., 2022), we demonstrate in the present study that astrocytes may represent an important target for KD action. Specifically, the severe astrogliosis seen in a clinically relevant murine model of developmental epilepsy can be mitigated by KD treatment, in parallel with a significant reduction in SRS. These actions are likely mediated through multiple mechanisms and molecular targets. Furthermore, the reduced astrocytic hypertrophy in hippocampus induced by the KD is accompanied by restoration of the passive membrane conductance likely mediated by potassium channels. Our observations are novel with respect to KD action and are in line with the growing evidence that astrogliosis is an important pathophysiological factor in seizure genesis and epileptogenesis (Çarçak et al., 2023; Fedele et al., 2005). Indeed, restoration of normal astrocytic function increasingly represents a viable strategy for epilepsy therapeutics.

### The KD Improves Spatial Potassium Buffering

The extracellular potassium concentration ([K^+^]_out_) in the brain rises in response to conditions such as hypoxia-ischemia, hypoglycemia, and epilepsy, leading to significant disruptions in brain function. Astrocytes play a crucial role in maintaining potassium (K^+^) balance in the brain through a mechanism known as spatial potassium buffering. In addition, astrocytes are involved in other important functions such as glutamate uptake, water homeostasis, and facilitating synaptogenesis (Belov Kirdajova et al., 2020; Murakami & Kurachi, 2016). In this context, of obvious importance is the role that potassium channels play, not only in repolarization of the neuronal cell membrane during action potential firing, but also in astrocyte physiology.

Neuronal activity within the physiological range causes a local increase in extracellular potassium concentration ([K^+^]_o_) of less than 1 mM. However, during seizure activity, ([K^+^]_o_) can rise to 10–12 mM, reaching a plateau level and inducing significant effects of synaptic transmission and plasticity. Astrogliosis, the reactive response of astrocytes to injury or disease, often involves changes in the expression and function of various ion channels. In astrocytes, it has been reported that the two-pore domain TWIK-1/TREK-1 potassium channel heterodimers can mediate astrocytic passive conductance and cannabinoid-induced glutamate release from astrocytes (Mi Hwang et al., 2014). Given that our results did not directly implicate TREK-1 and TWIK-1 channels (Figure S1E), we turned our attention to Kir4.1 channels to determine whether the KD and/or BHB might affect either Kir4.1 expression and/or function.

Kir4.1 is a weakly inwardly rectifying potassium channel that is exclusively expressed in glial cells in the nervous system, with the highest levels found in astrocytes. Within astrocytes, Kir4.1 is implicated in several critical functions, including potassium ion homeostasis, maintaining the astrocyte resting membrane potential, contributing to high potassium conductance, regulating astrocyte cell volume, and facilitating glutamate uptake (Kinboshi et al., 2020; Olsen et al., 2015). Multiple research groups have demonstrated that inhibition of the *Kir4.1* channel leads to elevated extracellular potassium concentrations and compromised glutamate uptake (Imbrici et al., 2013). This elevation of glutamate levels in the synaptic cleft disrupts normal synaptic transmission and network-level communication (Wilcock et al., 2009). Downregulation of *Kir4.1* has also been linked to disorders characterized in part or chiefly by neuroinflammation such as multiple sclerosis and Alzheimer’s disease (Milton & Smith, 2018; Obara-Michlewska et al., 2015; Zhang et al., 2017).

There is substantial evidence indicating that Kir4.1 is the primary channel involved in astrocyte-mediated spatial buffering and may account for up to 45% of potassium buffering in the hippocampus (Brill et al., 2015; Morris et al., 2020). Hence, pathologic alterations in Kir4.1 expression and activity would be expected to modulate potassium buffering and extracellular potassium homeostasis to promote neuronal hyperexcitability, a hallmark feature of epilepsy (Brill et al., 2015; Huffels et al., 2022; Marnetto et al., 2017; Srivastava et al., 2012).

Various research groups, including our own, have previously demonstrated that KB can influence hippocampal neuronal activity through Kir6.2 (K_ATP_) channels (Kim et al., 2015; W. Ma et al., 2007; Rho & Boison, 2022; Stafstrom & Rho, 2012). While we were not able to demonstrate significant effects of BHB in modulating potassium currents in Kir4.1-transfected HEK cells, we did observe a marginal increase after bath application.

### The KD Evokes Restoration of Differential Gene Expression

Prior investigations have examined gene expression changes evoked by KD therapy in various model systems, and the results suggest not only potential therapeutic targets for certain disorders but also highlight an overall homeostatic effect that metabolism-based therapies might afford (Jiang et al., 2022; Mychasiuk & Rho, 2017; Stafford et al., 2010). Here, we observed from our RNA-seq data a prominent normalizing effect of the KD in our animal model of epilepsy (Figure 6). Among the 332 differentially expressed (DE) genes identified between WT and KO mice, a substantial majority (300) of these were no longer differentially expressed after KD treatment and a subset of which are involved in the pathophysiology of astrogliosis (Figure 2E). The changes in gene expression profiles were also found to correlate with results obtained from our immunocytochemical and quantitative PCR (qPCR) analysis as demonstrated in Figure 3. Interestingly, in WT mice fed a KD, there were minimal changes in gene expression patterns, suggesting that the KD exerts disease-specific effects and that prior findings of diet-induced alterations in normal brain may not be relevant to its therapeutic underpinnings.

A recent transcriptomics study identified six essential genes whose expression was altered by the KD, notably an increased expression of ADCY3 (Lin et al., 2024). These authors demonstrated that activation of the ADCY3-initiated cAMP signaling pathway enhanced neuronal inhibition. Another study implicated the neuregulin 1/ErbB4 signaling pathway in elevating levels of γ-aminobutyric acid (GABA) in the brain (Wang et al., 2021). Both these mechanisms would be expected to contribute to the anti-seizure effects of the KD. And in further support of our findings, a recent single nucleus RNA-seq (snRNA-seq) study in a middle cerebral artery occlusion model of ischemic stroke and reperfusion injury revealed heterogeneity of gene expression changes in reactive astrocytes, with substantial overlap with the differentially expressed genes in our KO model (H. Ma et al., 2022).

Finally, in yet another report, the KD was shown to improve motor function in a mouse spinal cord injury (SCI) model, with gene expression changes similar to what we found in our epileptic KO mice (Streijger et al., 2013). It is noteworthy that after SCI, astrogliosis scar formation occurs and the heterogeneous astrocyte populations play important roles in axon regeneration and recovery from injury (Anderson et al., 2016; Escartin et al., 2021; Silver & Miller, 2004). As we observed that the KD can reverse some of the same gene expression changes seen in this SCI study (Figure 6), our data provide further evidence that metabolic therapies such as the KD could be a therapeutic option for patients with SCI (Demirel et al., 2020; Streijger et al., 2013; Tan et al., 2020).

Despite our current findings, it is emphasized that there are limitations to what can be interpreted from bulk RNA-seq experiments. First and foremost, there is a lack of cell-type specificity, and it is unknown whether any of the expression changes seen between WT and KO mice, with either a SD or KD, can be explained by alterations in neuronal (not necessarily astrocytic) gene expression. Furthermore, it is not possible to determine whether important cell-specific changes in relatively rare cell populations are masked when gene expression changes are averaged across the tissue sample. Yet, despite this well-recognized limitation, and other potentially confounding variables, our initial results suggest that the KD can restore – at least in part – pathophysiological changes at the level of astrocytes in the epileptic brain.

### Summary

To our knowledge, this is the first report that the KD can mitigate aberrant astrogliosis in the epileptic brain, and as such restore astrocyte passive membrane conductance and normal potassium homeostasis, possibly through the action of Kir4.1 channels and through reversing gene expression changes associated with astrogliosis. And although there are several studies linking astrocytic ion channels to their function in the context of a KD, our study is the first to directly demonstrate the relevance of these inter-relationships in an animal model of epilepsy (Çarçak et al., 2023; Ohno, 2018; Purnell et al., 2023). Collectively, our data strengthen the growing evidence for the critical role astrocytes play in epilepsy (and potentially other neurological disorders) and a therapeutic mechanistic target for KD action.

## Methods

### Animals and Feeding

Wild-type (WT) and *Kcna1*-null (KO) mice on a C3HeB/FeJ background strain were bred from cryo-recovered heterozygous animals (Jackson Laboratories, Bar Harbor, ME) and weaned on postnatal day 20 (P20). Mice were maintained on a 12-hr light/dark cycle with lights on at 7:00 a.m. in a temperature-controlled room. Genotyping was outsourced to Transnetyx. Efforts were made to minimize pain and the total number of animals required. All experimental procedures were approved by the Institutional Animal Care and Use Committee (IACUC) at the University of California San Diego (protocol number S09186) and conformed to the National Institutes of Health guidelines on the care of laboratory animals. Upon weaning, mice were separated by gender, housed in standard cages, and placed ad libitum on either a standard diet (SD) or KD (F3666; BioServ, Frenchtown, NJ). Blood BHB and glucose levels were assayed with KetoMojo^©^ (Napa, California and determined at multiple time-points during KD and SD treatment.

### Video-EEG

A prefabricated head-mount was surgically implanted in 3-week-old KO mice, and continuous video-EEG monitoring was initiated 2 days later for 14 days using a Pinnacle tethered system (Lawrence, Kansas, USA). The KD was administered *ad libitum* to mice in the experimental treatment group (n=5). The KD was placed in 3.3 cm petri dishes and replaced daily. The control group of mice (n=5) was given a SD and similarly underwent video-EEG monitoring. Seizures were monitored and behaviorally scored using the Racine scale as previously described (K. A. Simeone et al., 2016). Specifically, stage 1, myoclonic jerk; stage 2, head stereotypy; stage 3, limb clonus, tail extension, and single rearing event; stage 5, continuous rearing and falling; and stage 5, severe tonic-clonic seizures. A seizure event was only counted if: 1) it could be classified as any of the stages above; and 2) it was accompanied with single or sustained EEG bursts. Gamma power was calculated from the EEG recordings from both the KD- and SD-fed mice. Six hours of spontaneous EEG recordings were analyzed when the animal was active (ZT15 – ZT21). Power spectral density (PSD) of the EEG signal was calculated using Welch’s method implemented in MATLAB.

### RNA Sequencing and Analysis

The library preparation and the bulk RNA sequencing was performed in mouse hippocampus from four groups of animals (WT/SD, WT/KD, KO/SD,KO/KD) through Azenta Life Sciences (La Jolla, California USA). The bulk RNA-seq analysis was performed using DESeq2, Gene Ontology and network analysis. Using DESeq2, we found differentially expressed (DE) genes between the four different groups (WT/SD, WT/KD, KO/SD, KO/KD). For 4-way analysis, we compared the DE genes between WT and KO under two conditions. We performed Gene Ontology (GO) gene enrichment analysis with the genes that are significantly changed (from WT to KO) in SD but not in KD. For recovery analysis, we specifically focused on the significant DE genes induced by *Kcna1* deletion, and significant DE genes but reversely by the KD. Most heatmaps were created using the heatmap3 function.

### Immunohistochemistry

Mice from each of the four groups (WT/SD, WT/KD, KO/SD, KO/KD) were transcardially perfused under isoflurane anesthesia or after intraperitoneal injections of ketamine-xylazine at 0.015 ml/ gram of body weight. Perfusion was conducted with phosphate-buffered saline, followed by 4% paraformaldehyde (PFA) solution before the brains were extracted, fixed in a 4% PFA solution 4°C overnight and cryoprotected in a 30% sucrose solution for 2 to 3 days. 40 μm coronal 477 frozen sections were prepared using a cryostat (Leica CM1860) and 340μm coronal sections 478 were cut using a vibratome (Leica, VT1000S).

Fluorescence immunohistochemistry was performed on the (40μm & 340μm) coronal sections. Each section was washed three times for 5 minutes using 1 x tris-buffered saline (TBS) solution. After washing, coronal sections were permeabilized in 0.2% Triton X-100 in 1x TBS containing 0.5 % Tween20 for 1 hour and washed 3 times for 5 minutes each using 1 x TBS solution. Next, the slices were blocked in 0.2% Triton X-100 in 1 x TBS containing 3% normal serum (donkey/goat: sigma Aldrich, D9663, G9023) for 1 hour. After blocking, sections were incubated with the primary antibodies to GFAP (mouse antibody,1:1000, Abcam, Anti-GFAP antibody #ab7260, and Kir 4.1 (rabbit antibody, 1:200, alomon labs, #APC-035) at 4°C for 72 hours (3 days). Sections were washed 3 times for 10 minutes each using 1 x TBS. Subsequently, secondary antibodies tagged with anti-rabbit Alexa Fluor568 (red fluorescence, 1:250), anti-mouse Alexa Fluor488 (green fluorescence, 1:250) were used, and slices were incubated for 24 hours. Then, 4′,6-diamidino-2-phenylindole (DAPI, 1:1000, Thermo Scientific™:DAPI and Hoechst Nucleic Acid Stains #62248) was used to label nuclei and incubated for 25 minutes. Stained sections were examined using confocal laser scanning microscope (Zeiss LSM 880 Confocal with Airyscan)

### Electrophysiology

#### Preparation of acute hippocampal slices and Sulforhodamine 101(SR101) staining

Acute coronal 340 μm-thick hippocampal brain slices were prepared from KO or WT mice on either a SD or a KD using a vibratome (Leica, VT1000S) in ice-cold oxygenated artificial cerebrospinal fluid containing (in mM) 130 NaCl, 2.5 KCl, 1.25 KH_2_PO_4_, 3.0 MgCl_2_, 1.0 CaCl_2_, 26 NaHCO_3_, and 10.0 D-glucose. Slices were then transferred to a beaker with a holding basket containing 0.6 μM Sulforhodamine 101 (SR101) in aCSF at 34°C for 30 min until used for experiments. The hippocampal slices were visualized using a Zeiss Axioskop 2 FS plus microscope equipped with epifluorescence.

#### For Hippocampal astrocyte whole-cell patch recording

The SR101-positive cells were selected for whole-cell patch clamp recording. The slices were stored at RT until the experiments were performed. Patch pipettes had a resistance of 3.5–4.5 MΩ when filled with pipette solution containing (in mM) 140 KCl, 10 HEPES, 5 EGTA, 2 Mg- ATP, and 0.2 Na-GTP, adjusted to pH 7.4 with KOH. Whole-cell patch recordings were performed on hippocampal astrocytes with a voltage-clamp configuration using an Axopatch 700B (Axon Instruments, Union City, CA, USA). Whole-cell membrane currents were amplified by the Axopatch 200A. Currents were elicited by 1-s ramps descending from +50 mV to −150 mV (from a holding potential of −70 mV). Data acquisition was controlled by pCLAMP 10.2 software (Molecular Devices, Sunnyvale, CA, USA). The Digidata 1440A interface was used to convert digital–analog signals between the amplifier and the computer. Data were sampled at 5 kHz and filtered at 2 kHz. Cell membrane capacitance was measured using the ‘membrane test’ protocol built into pClampex.

#### Electrophysiological recording in CA1 pyramidal neuron

The pipette solution contained (in mM) 120 potassium gluconate, 10 KCl, 1 MgCl_2_, 0.5 EGTA, and 40 HEPES (with the pH adjusted to 7.2 with KOH). Recordings were considered stable when the series and input resistance and the RMP did not vary by >20%. Recordings were filtered at 2 kHz and digitized at 10 kHz. In the current clamp experiments, a current was applied at increasing stepwise increments of 5 pA, each with a duration of 1.2 s. The data were collected with a MultiClamp 700B amplifier (Molecular Devices) using Clampex10 acquisition software (Molecular Devices) and digitized with a Digidata 1440A (Molecular Devices). Whole-cell patch-clamp recordings were performed using a MultiClamp 700A amplifier and pClamp 9.2 software (Molecular Devices). A minimum of 1 GΩ seal resistance was required before rupturing the membrane into a whole-cell configuration. The membrane potential (VM) was read either in “I = 0” mode or measured directly in current clamp mode without applying any holding currents. For neuronal recording, the access resistance (Ra), membrane resistance (RM), and membrane capacitance (CM) were measured from the “Membrane test” protocol built into the pClampex9.2. For quality control, a Ra < 20 MΩ is used to include neurons in the final data analysis.

#### Electrophysiological recording in transfected Kir4.1-GFP, Control-GFP HEK293T cells

Whole-cell membrane currents were amplified using the Axopatch 200A patch clamp system. Currents were elicited by 1-s ramps descending from +50 mV to −150 mV (from a holding potential of −70 mV). Data acquisition was controlled by pCLAMP 10.2 software (Molecular Devices). A Digidata 1440A interface was used to convert digital–analog signals between the amplifier and the computer. Data were sampled at 5 kHz and filtered at 2 kHz

### Quantitative Real-Time Polymerase Chain Reaction (qPCR)

Samples for qPCR were acquired from WT and KO mice at postnatal age 42 (P42). Half of the hippocampus (with the other half being used for Western blotting) was homogenized in 700 µL of Trizol, and 200 µL of chloroform was added. Isopropanol was then used to precipitate out the RNA, which was subsequently washed with 70% EtOH, and solubilized in ultra-pure RNAse- free water. To synthesize cDNA, an Applied Biosystems High-Capacity cDNA Reverse Transcription Kit was used to make up a total of 10 µL of master mix and 10 µL of RNA working solution. The tubes were sealed and subjected to the following thermal protocol in a thermal cycler: 25°C, 10 min; 37°C, 120 min; 85°C, 5 min; 4°C, hold. Next, the cDNA was diluted in RNase-free water at a 1:15 dilution, and the remaining cDNA was stored at -80°C. For qPCR, a mix was prepared using 10 µL of 2X TaqMan PreAmp Master Mix, 1 µL of 20X primer (to make a total of 11 µL) and 9 µL of diluted cDNA. The plate was then sealed, and qPCR was run using a QuantStudio 3 Real-Time PCR System. Using C(t) values from qPCR using specific primers, we calculated A.U. values of each protein, GFAP, Kir4.1, TWIK-1 and TREK-1. To normalize the expression based on the astrocyte region, we calculated the volume by taking the square root of GFAP surface area and applying the cubic. We divided each A.U. value with the calculated volumes to get expression estimates. Using C(t) values from qPCR using specific primers, we calculated A.U. values of each protein, GFAP, Kir4.1, TWIK-1 and TREK-1. To normalize the expression based on the astrocyte region, we calculated the volume by square root of GFAP surface area and applied the cubic. We divided each value of A.U. value with the calculated volume values to get expression estimates.

### Western Blotting

Samples for Western blotting were acquired from WT and KO mice at P42. The remaining half of the hippocampus (with the other half being used for qPCR) was homogenized in 300 µL of lysis buffer and 6 µL of 50X of protease inhibitor. The lysis buffer consisted of: 0.5% SDS, 0.05M Tris-Cl, and 1 mM dithiothreitol (pH 8.0). Samples were incubated with a sodium dodecyl sulfate-polyacrylamide gel electrophoresis (SDS-PAGE) sample buffer for 5 min at 95°C. Each sample (10 µg/lane) was run on a 10% Bis-Tris gel for 1 hour and then transferred for 60 min to a PVDF membrane. The membrane was then incubated in a blocking solution containing 0.5% milk powder in Tris Buffered Saline with 0.1% Tween20 for 1 hr, and overnight with the corresponding antibodies. The next day, the blots were incubated for 60 minutes with an anti-rabbit or anti-mouse IgG antibody. Blots were then developed using a chemiluminescence method (using Genesee Scientific Prosignal Femto ECL Reagent).

### Cell Culture and Transfection

For cell cultures, HEK293 cells were cultured in high glucose Dulbecco’s modified Eagle’s medium (DMEM: Sigma Aldrich) which was supplemented with 10% Fetal Bovine Serum (FBS: Gibco). The cells were cultured in a 24-well plate on Poly-D-Lysine coated coverslips, at 50% confluency with 1 mL medium in each well, and were placed in a CO_2_ incubator overnight. Plasmid DNA and 1 mg/ml PEI in a 1:3 ratio was mixed with 200 μL of DMEM, and the mixture was incubated for 30 minutes at RTP. Prior to adding the transfection media to the cells, the media was changed, and the mixture was dropped into each well and incubated overnight, with a media change after 17 hours and every other day after that.

### Comparative Analysis with Public Dataset

We downloaded the original differential expression dataset table from Burda et al. 2022. (http://tr.astrocytereactivity.com). We compared our DE genes log2 Fold Change with these datasets for the analysis presented in Figure 6.

### Statistics/statistical design

All experiments were designed with sufficient animals/groups to ensure adequate statistical power on pilot experiments. All data are presented as means ± standard error of the mean (S.E.M). The significance of data for comparison was assessed by Student’s t-test (paired t-test) or one-way ANOVA followed by Turkey’s post hoc test, and significance levels are given as: n.s: not significant, * p < 0.05, ** p < 0.01, *** p < 0.001, and **** p < 0.0001.The statistical analysis was carried out using Prism 9.0 software (GraphPad Software, San Diego, CA, USA). For RNA-seq analysis, we employed size factor normalization for comparison.

## Data Availability

The sequencing data is available at GSE264537.

## Acknowledgement

This work was conducted at the University of California San Diego (UCSD) and supported by start-up funding provided to J.M.R. by the Rady Children’s Hospital and the Department of Neurosciences at UCSD. 619

Current address of the first author: Jae Hyouk Choi, Burke Neurological Institute, Weill Cornell 620 Medicine, 785 Mamaroneck Avenue, White Plains, NY 10605, USA. Email: 621 The authors thank the UCSD for technical assistance and collaborators for their intellectual contributions. We also acknowledge the Burke Neurological Institute and Weill Cornell Medicine for their ongoing institutional support. Finally, we extend our gratitude to the study participants and lab members for their invaluable contributions.

---

| Gene | Standard Diet WT VS KO |  |  | Ketogenic Diet WT VS KO |  |  |
| --- | --- | --- | --- | --- | --- | --- |
|  | Log 2 Foldchange | Adjusted p-value | Significance | Log 2 Foldchange | Adjusted p-value | Significance |
| Aspg | -1.245828047 | 0.003146549 | *** | -1.096198574 | 0.996466074 | ns |
| SLC25A18 | 0.428599965 | 0.0032651 | *** | 0.042843265 | 0.996466074 | ns |
| RIMS1 | 0.396426433 | 0.005159348 | *** | 0.02512343 | 0.996466074 | ns |
| ACAP3 | 0.3318534 | 0.013083462 | *** | 0.146938492 | 0.996466074 | ns |
| RASSF8 | -0.55696353 | 0.013083462 | *** | -0.513885298 | 0.372768099 | ns |
| HES1 | 0.680796502 | 0.017367813 | *** | -0.280964425 | 0.996466074 | ns |
| SESN1 | -0.382121323 | 0.018170978 | *** | -0.071304455 | 0.996466074 | ns |
| ETNPPL | -1.535105173 | 0.018947103 | *** | -0.388805161 | 0.996466074 | ns |
| Junb | 0.512060224 | 0.019794159 | *** | 0.671589282 | 0.060324729 | ns |
| LINGO1 | 0.43291446 | 0.021542434 | *** | -0.254402792 | 0.996466074 | ns |
| Sparc | -0.712175602 | 0.021542434 | *** | -0.28713871 | 0.63727772 | ns |
| Jun | -0.392517907 | 0.02254442 | *** | -0.29373586 | 0.081682662 | ns |
| Spp1 | -4.569174281 | 0.022619205 | *** | -0.485649124 | 0.996466074 | ns |
| Cd44 | -1.107167841 | 0.02267334 | *** | -1.993092367 | >1 | ns |
| Ifit3 | -1.123225319 | 0.027191497 | *** | -0.417199692 | 0.996466074 | ns |
| Clu | -0.414571426 | 0.027466437 | *** | 0.05017985 | 0.996466074 | ns |
| COL5A3 | -0.968215297 | 0.028689436 | *** | -1.250572527 | 0.820065546 | ns |

