## Supplementary figures and images for "A ketogenic diet mitigates hippocampal astrogliosis in epileptic brain"

### Figure S1

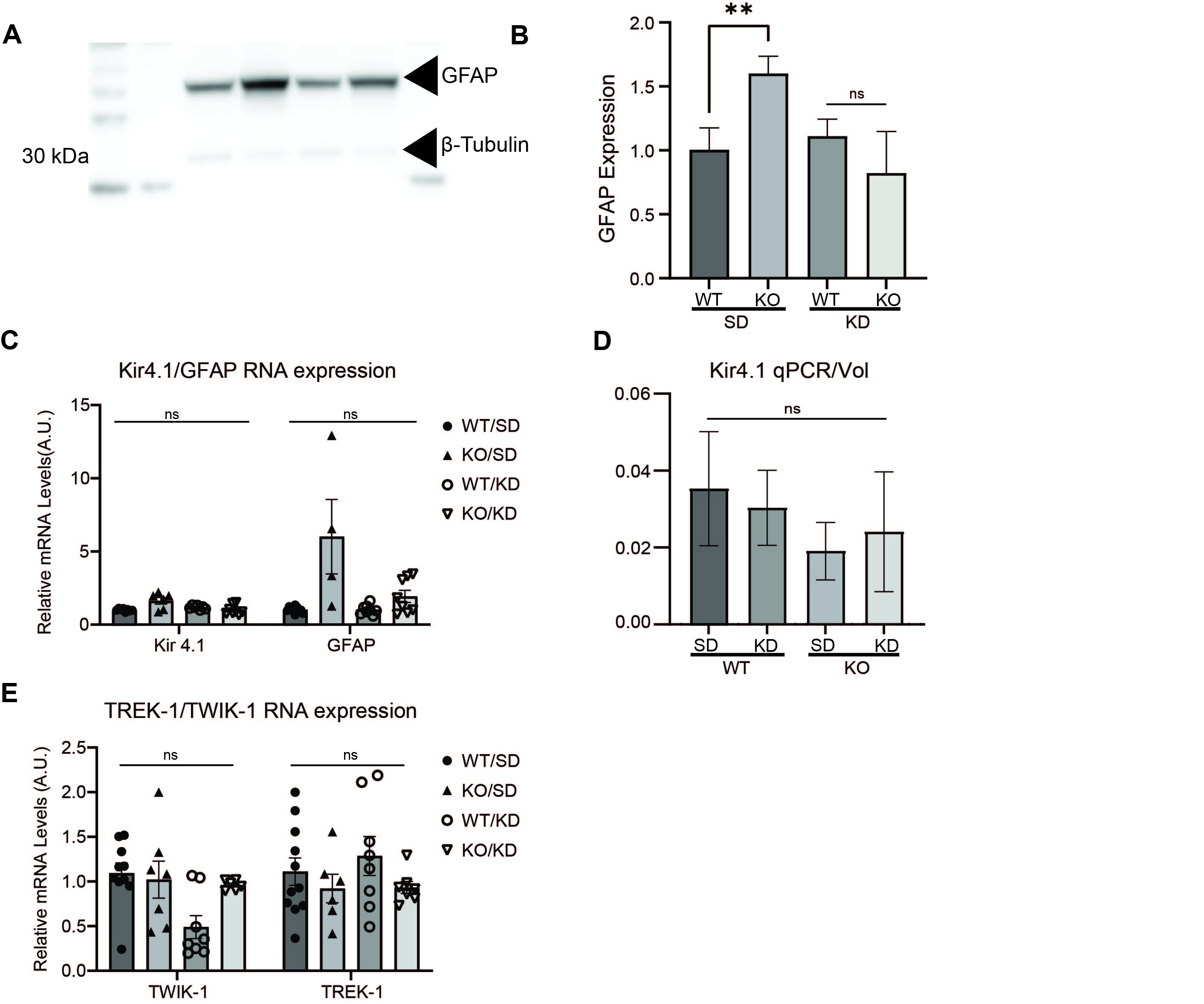

### Figure S2

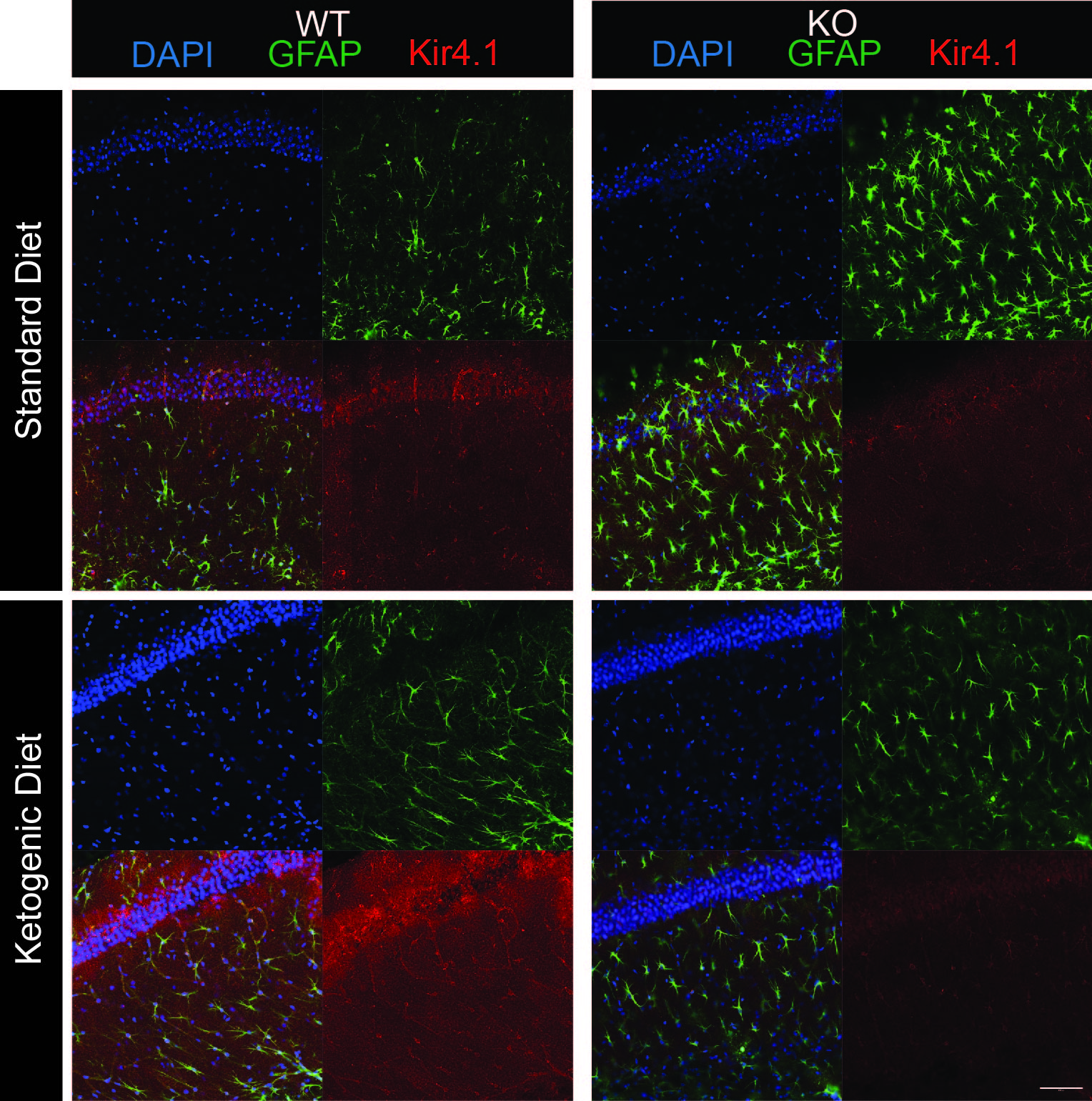

### Figure S3

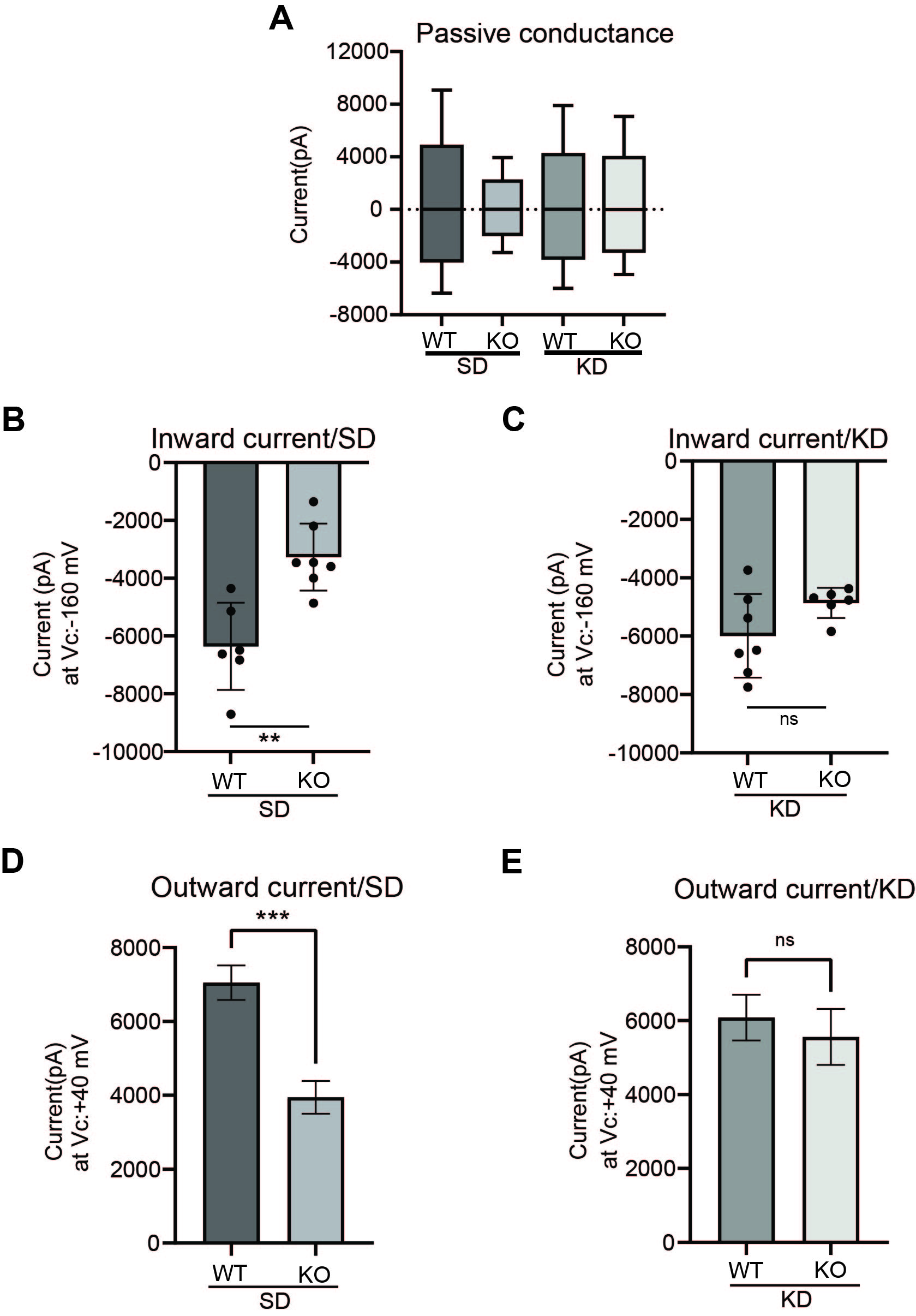

### Figure S4

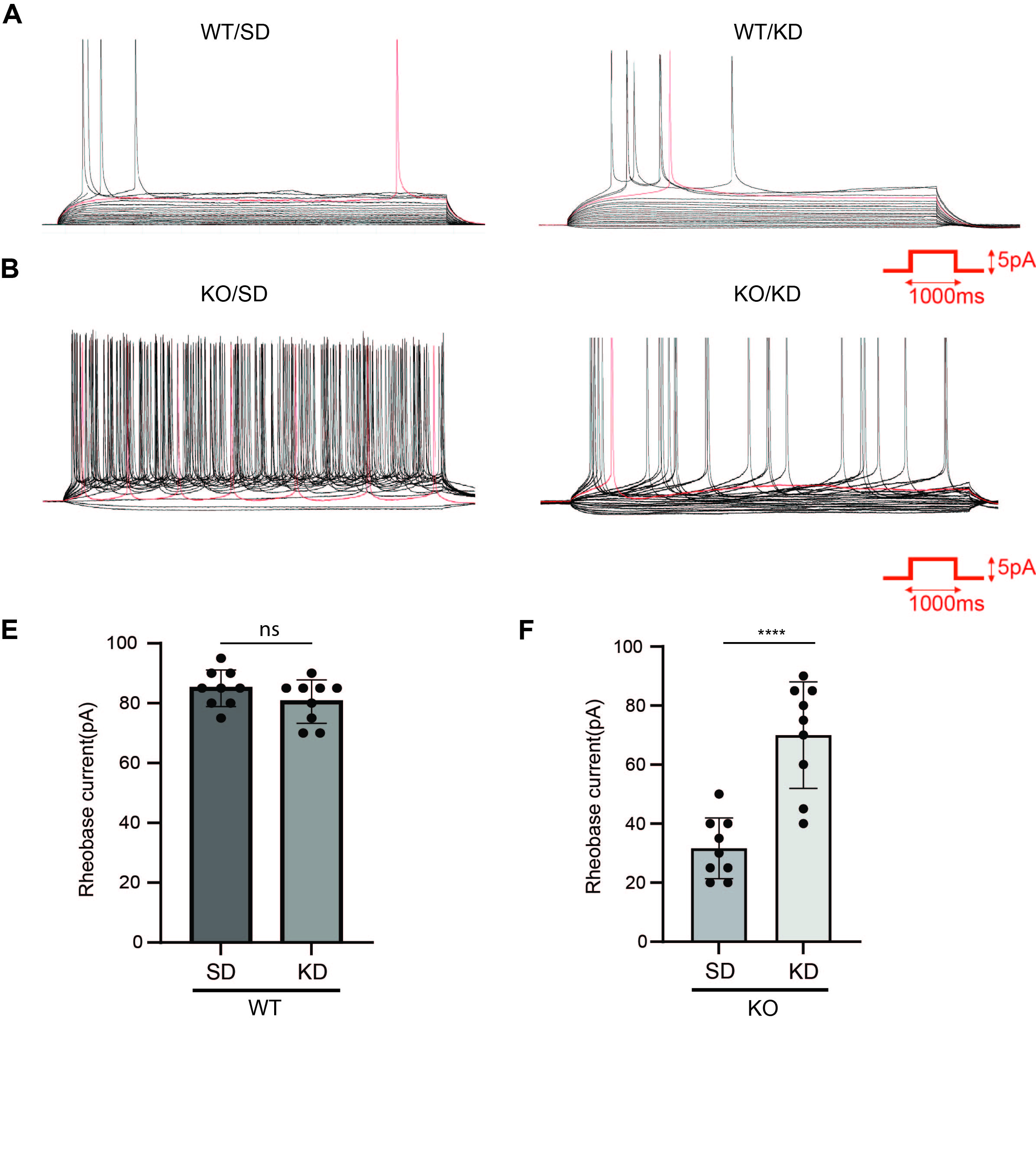
